# EcOH: *In silico* serotyping of *E. coli* from short read data

**DOI:** 10.1101/032151

**Authors:** Danielle J. Ingle, Mary Valcanis, Alex Kuzevski, Marija Tauschek, Michael Inouye, Tim Stinear, Myron M. Levine, Roy M. Robins-Browne, Kathryn E. Holt

## Abstract

The lipopolysaccharide (O) and flagellar (H) surface antigens of *Escherichia coli* are targets for serotyping that have traditionally been used to identify pathogenic lineages of *E. coli*. As serotyping has several limitations, public health reference laboratories are increasingly moving towards whole genome sequencing (WGS) for the rapid characterisation of bacterial isolates. Here we present a method to rapidly and accurately serotype *E. coli* isolates from raw, short read sequence data, leveraging the known genetic basis for the biosynthesis of O- and H-antigens. Our approach bypasses the need for *de novo* genome assembly by directly screening WGS reads against a curated database of alleles linked to known *E. coli* O-groups and H-types (the EcOH database) using the software package SRST2. We validated our approach by comparing *in silico* results with those obtained via serological phenotyping of 197 enteropathogenic (EPEC) isolates. We also demonstrated the utility of our method to characterise enterotoxigenic *E. coli* (ETEC) and the uropathogenic *E. coli* (UPEC) epidemic clone ST131, and for *in silico* serotyping of foodborne outbreak-related isolates in the public GenomeTrakr database.

## Introduction

Differentiation of isolates of *Escherichia coli* is commonly performed by serological typing (serotyping) of the highly polymorphic somatic- (O) and flagellar- (H) antigens (DebRoy *et al*., 2011; Wang *et al*., 2003). The O-antigen is an integral part of the lipopolysaccharide (LPS) in the outer membrane of Gram-negative bacteria, whilst the H-antigen projects beyond the cell wall and provides cell motility (Li *et al*., 2010; Wang *et al*., 2003). Currently there are 182 *E. coli* O-groups and 53 H-types recognised by serotyping (Croxen *et al*., 2013; Iguchi *et al*., 2014; Joensen *et al*., 2015). Serotyping involves performing a series of agglutination reactions with panels of antisera, and is expensive in terms of both labour and reagent costs (Achtman *et al*., 2012; Fratamico *et al*., 2009). In addition, the interpretation of these assays is subjective and relies on antisera that vary in titre and specificity according to the source and integrity of the serum. Further, a significant proportion of *E. coli* isolates (approximately one quarter) are serologically ‘untypeable’ due to cross-reactivity or a lack of reaction with available antisera (DebRoy *et al*., 2011). For these reasons there has been a shift away from serological phenotyping of *E. coli*, towards inference of O- and H- genotypes using molecular methods (Jenkins, 2015).

O-antigen biosynthesis in *E. coli* is encoded in gene clusters that are typically located between the chromosomal housekeeping genes *galF* and *gnd/ugd* (Iguchi *et al*., 2014; Liu *et al*., 1996). The genes required to synthesise this antigen fall into three classes: (i) sugar synthesis genes, (ii) glycosyl transferase genes, and (iii) O-antigen processing genes (Samuel & Reeves, 2003). Two distinct O-antigen pathways are known: (i) the Wzx/Wzy-dependent pathway, encoded by the *wzx* (O-antigen flippase) and *wzy* (O-antigen polymerase) genes, and (ii) the ABC transporter pathway, encoded by *wzm* and *wzt* (Feng *et al*., 2004; Samuel & Reeves, 2003). In general, variation in these gene sequences correlates with structural variation in the carbohydrate residues that make up each O-antigen (DebRoy *et al*., 2011; Wang *et al*., 2003). Because of this, the sequences of these genes can be used for O-antigen genotyping (Joensen *et al*., 2015; Mentzer *et al*., 2014). Nevertheless, genotype-phenotype relationships for some O-groups are unexpectedly complex. For example, two distinct gene clusters are associated with the same O45 serotype (Plainvert *et al*., 2007), whereas some distinct O-antigens are encoded by near-identical gene clusters (Iguchi *et al*., 2014).

H-antigen specificity is determined by flagellin, which is the protein subunit of flagella. This protein is encoded by *fliC* in 43 of the 53 serologically defined H-types (Wang *et al*., 2003). PCR detection of *fliC* alleles has been used for molecular H-typing for some time (Wang *et al*., 2003). However, some *E. coli* isolates have an alternative flagellar phase, due to the presence of an additional flagellin gene *(flnaA, fllA, fmlA* or *flkA)*, similar to those found in *Salmonella* species (Feng *et al*., 2008; Ratiner, 1998; Ratiner *et al*., 2010; Tominaga, 2004; Tominaga & Kutsukake, 2007).

As the cost of high-throughput short read DNA sequencing declines, public health laboratories are increasingly moving away from phenotyping and towards whole genome sequence (WGS) based typing of bacteria including *E. coli* (Joensen *et al*., 2015; Kwong *et al*., 2015). Given the strong genetic basis for O- and H-antigenic variation in *E. coli*, the availability of genome data provides a valuable opportunity to infer serotypes at little or no additional cost. Here we present a method to rapidly and accurately serotype *E. coli* isolates from raw, short read sequence data, by screening reads directly against a curated database of alleles linked to known *E. coli* O-groups and H-types (the EcOH database, presented here) using the software package SRST2 (Inouye *et al*., 2014). The EcOH database can also be used to infer serotypes from assembled genome data using BLAST or other sequence comparison tools, which will become increasingly useful as long-read sequence data become more common. We validated our approach by comparing *in silico* predicted serotypes to those determined phenotypically in a public health reference laboratory, and demonstrated the utility of *in silico* serotyping to characterise more than 1,000 *E. coli* isolates including enteropathogenic *E. coli* (EPEC), enterotoxigenic *E. coli* (ETEC), the uropathogenic *E. coli* (UPEC) clone ST131 and foodborne outbreak-associated isolates of *E. coli* deposited in the public GenomeTrakr database.

## Methods

### Curation of the EcOH database

The EcOH database of O- and H-type encoding sequences was initially constructed in 2014 from publically available sequences identified in GenBank by reviewing the literature on the PCR detection of *E. coli* O- and H-types (DebRoy *et al*., 2011; Ratiner *et al*., 2010; Wang *et al*., 2003). This was updated by a further review in May 2015 (Iguchi *et al*., 2014; Joensen *et al*., 2015). Twelve novel O loci identified in the present study were also included. The resulting EcOH database includes sequences of alleles for *wzm* and *wzt*, or *wzx* and *wzy*, covering 180 O-groups; and *fliC, flnA, fmlA, flkA* and *fllA* allele sequences covering all 53 known H-types. Details of all sequences in the EcOH database are provided in **Table S1**. The EcOH database is available at https://github.com/katholt/srst2.

### Publically available sequence data used in this study

Details of the short read Illumina data used in this study are provided in **Table 1**. A total of 41 complete *E. coli* genomes were downloaded from PATRIC (Wattam *et al*., 2013), with the accession numbers given in **Table S2**. Serologically determined O-groups were identified in the GenBank entries or associated literature for 40 of these genomes (and H-types for 20) (**Table S2**). The Achtman multi-locus sequence typing (MLST) scheme for *E. coli* (Wirth *et al*., 2006), now hosted at Warwick University (http://mlst.warwick.ac.uk/mlst/dbs/Ecoli), was downloaded using the getmlst.py script included in the SRST2 package (https://github.com/katholt/srst2). An SRST2-formatted version of the ARG-ANNOT antimicrobial resistance gene database (Gupta *et al*., 2013) was downloaded from https://github.com/katholt/srst2.

**Table 1.**
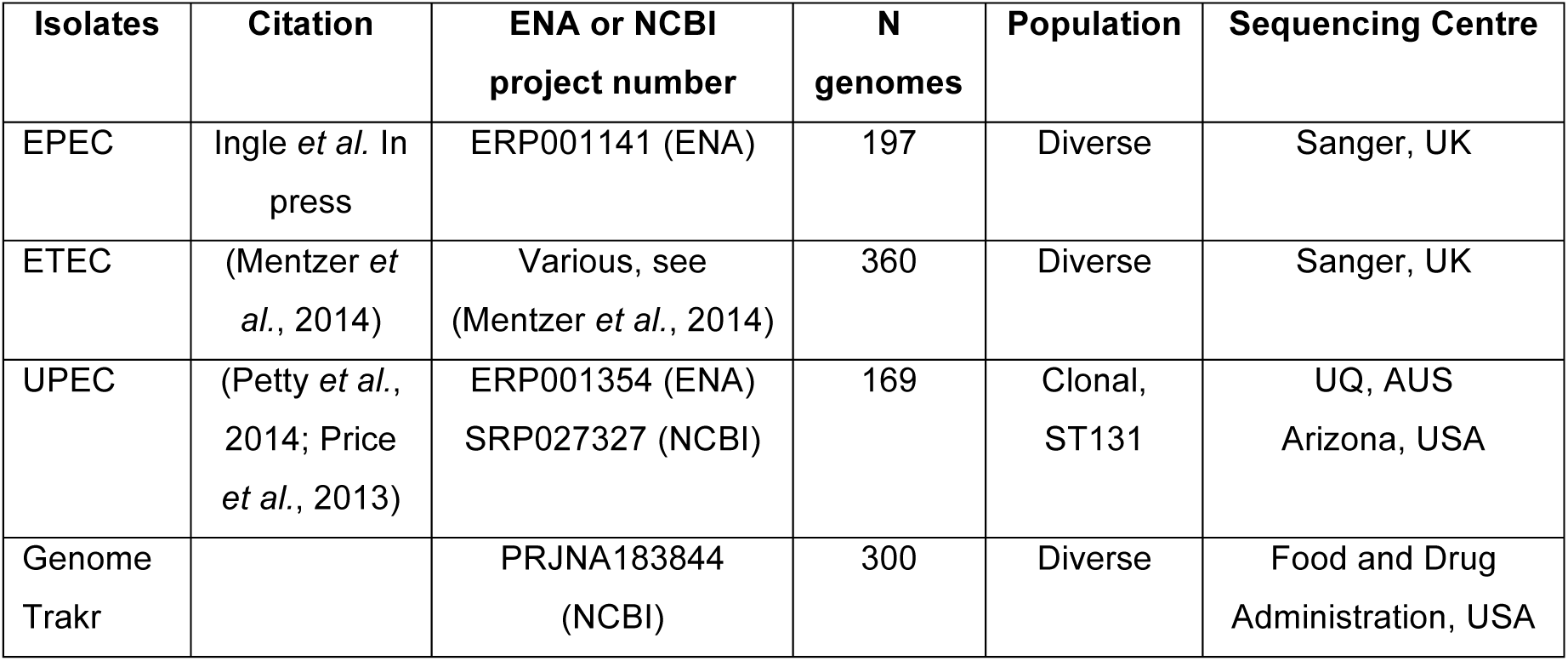
Datasets used to assess accuracy and utility of the EcOH database

| Isolates | Citation | ENA or NCBI project number | N genomes | Population | Sequencing Centre |
| --- | --- | --- | --- | --- | --- |
| EPEC | Ingle *et al.* In press | ERP001141 (ENA) | 197 | Diverse | Sanger, UK |
| ETEC | (Mentzer *et al.*, 2014) | Various, see (Mentzer *et al.*, 2014) | 360 | Diverse | Sanger, UK |
| UPEC | (Petty *et al.*, 2014; Price *et al.*, 2013) | ERP001354 (ENA)<br>SRP027327 (NCBI) | 169 | Clonal, ST131 | UQ, AUS<br>Arizona, USA |
| Genome Trakr |  | PRJNA183844 (NCBI) | 300 | Diverse | Food and Drug Administration, USA |

### Assembly and BLAST analysis

100 bp PE Illumina reads were generated previously for 197 EPEC isolates (Ingle **et al**, in press) and assembled using Velvet and Velvet Optimiser (Zerbino & Birney, 2008). Reads and assemblies are available in the European Nucleotide Archive (ENA) under ERP001141 (Ingle *et al*, in press). Here, we generated alternative assemblies using SPAdes (v3) (Bankevich *et al*., 2012) with error correction and kmer lengths of 21, 33, 55, 77 and 89. The resulting contigs were extended with the scaffolder SSPACE (Boetzer *et al*., 2011); gaps within the scaffolds were closed using GapFiller (Boetzer & Pirovano, 2012) and then further extended with AlignGraph (Bao *et al*., 2014).

Both sets of assemblies were screened against the EcOH database using BLAST+ *(blastn)*. A genotype call was made where a hit was identified with ≥90% coverage of a query sequence at ≥90% nucleotide identity. Note that as the SPAdes assemblies yielded fewer genomes with BLAST+ hits to O-antigen loci, these assemblies were discarded and all results were reported as comparisons of SRST2 data to assembly-based analysis using the Velvet Optimiser assemblies. This makes the comparison as generous as possible towards the competing method of assembly-based analysis.

### SRST2 analysis

SRST2 was run with default parameter settings, such that a genotype call reflects detection of reads covering ≥90% of the length of a query locus at ≥90% nucleotide identity. Where multiple alleles of the same locus appears in the database, SRST2 reports the best-scoring allele as the genotype call (Inouye *et al*., 2014). A confident genotype call is defined as one exceeding the minimum depth cut-offs (Inouye *et al*., 2014). Here we used the SRST2 default values of ≥5x mean read depth across the query locus to define a confident call.

### Phenotypic characterisation of 197 EPEC isolates

Isolates were subcultured on Luria-Bertani agar and incubated overnight at 37°C before being submitted to the National *E. coli* Reference Laboratory at the Microbiological Diagnostic Unit Public Health Laboratory (MDU PHL) in Melbourne, Australia, for serotyping.O-and H-serotyping utilised the standard tube agglutination tests, adapted for U-bottomed microtitre trays (Chandler & Bettelheim, 1974; Kauffmann, 1944).

### Characterisation of potential novel O-antigens

Where an O-group was determined via serological testing of an isolate, but no *wzx/wzy* or *wzm/wzt* genes were detected in the corresponding isolate’s genome, the *de novo* genome assembly was interrogated to identify potential novel O-antigen loci. For each such isolate the assembled contig containing the genes *galF* and *gnd*, which typically flank the O-antigen locus, was identified using BLAST and extracted using EMBOSS (Rice *et al*., 2000). The intervening sequences were annotated with Prokka (v1.11) (Seemann, 2014), using translated protein sequences from the EcOH database as the preliminary annotation source. We then used ACT (Carver *et al*., 2008) to visually compare the annotated sequences with full-length reference sequences for the corresponding O-group that had been identified by serology. Putative *wzx* and *wzy* alleles for these O-groups were identified based on (i) the annotation, (ii) sequence homology with the reference O-group sequences, and (iii) the presence of transmembrane domains identified using TMHMM (Krogh *et al*., 2001). These putative *wzx* and *wzy* gene sequences were added to the EcOH database with the suffix ‘var1, var2’, etc to differentiate them from the prototypical alleles (e.g. the novel *wzx* gene detected in isolates that were serologically phenotyped as O116 are labelled ‘wzx-O116var1’, whereas the prototypical O116 *wzx* gene is labelled wzx-O116, see **Table S1**).

### Analysis and visualisation of O- and H- antigen diversity and MLST data

For the EPEC and ETEC pathotypes, population structure was determined by constructing neighbour-joining trees based on Hamming distances between MLST allele profiles inferred from the genomes using SRST2. O- and H-types were plotted against these trees using R (plotting code is available in the *plotSRST2data.R* script within the SRST2 package at https://github.com/katholt/srst2). Diversity analyses, including Simpson index calculations and rarefaction plots, were performed using the *vegan* package for R (Oksanen *et al*., 2015).

### Analysis of ST131 UPEC

Illumina reads from a total of 169 isolates (accession numbers in **Table 1**) were mapped to the ST131 reference genome, SE15 (accession AP009378) (Toh *et al*., 2010) using the mapping-based pipeline RedDog (available at https://github.com/katholt/RedDog). Briefly, RedDog uses Bowtie2 (Langmead *et al*., 2009) to map short reads to the reference genome then uses SAMtools (Li *et al*., 2009) to call SNPs (Phred score ≥30, read depth ≥5x); consensus alleles at all SNP sites identified in the isolate collection are then extracted from each read set using SAMtools (Li *et al*., 2009) (Phred score ≥20 and unambiguous homozygous base call; otherwise allele call set to unknown ‘-’).

The resulting SNPs were filtered to include only those located within common genes (defined as genes with ≥95% coverage in ≥95% of the ST131 genomes analysed), yielding a total of 38,213 SNPs. The resulting SNP alignment was used as input to infer a maximum likelihood (ML) tree using RAxML (yielding 100% bootstrap support for all major nodes). The phylogeny was outgroup-rooted using the group comprising four closely related ST95 isolates (these had originally been identified as ST131 in PCR analysis for *rfb* and *pabB* genes, before MLST confirmed they were ST95 (Petty *et al*., 2014)).

### Analysis of GenomeTrakr

GenomeTrakr (NCBI BioProject: PRJNA183844) is a public repository of genome data from foodborne pathogens submitted by various laboratories including the US Food and Drug Administration and the Centers for Disease Control and Prevention. It includes raw Illumina reads and a kmer-based phylogeny of *E. coli* read sets, which is updated daily to incorporate newly submitted data. A subset of the most recently submitted read sets, together with the kmer tree, were downloaded from GenomeTrakr on 5 June, 2015. A subtree representing relationships between the 300 isolates was extracted from the full kmer tree, by removing all other tips from the tree using R packages *ape* (Paradis *et al*., 2004) and *Geiger* (Harmon *et al*., 2007).

## Results and Discussion

To identify unique sequences encoding *E. coli* O- and H- antigens, we began by curating a database (named EcOH) of 551 unique sequences representing known O- and H-types of *E. coli*, incorporating data from Iguchi *et al*. 2014 and several reviews (see **Methods**). Of the 182 currently recognised O-serogroups of *E. coli*, 180 corresponding genotypes were represented in the database by gene sequences for either *wzx* and *wzy*, or *wzm* and *wzt*. The two exceptions were O57 and O14, as isolates with these O-groups lack any of these genes and harbour only small O-antigen gene clusters, with no known polymerase or flippase genes, and only the housekeeping genes *galF, gnd* and *hisI* together with *ugd* and *wzz* which is not sufficient to delineate these O-groups. The EcOH database also includes sequences for all 53 known H-types, allowing for the detection of both *fliC* and *non-fliC* flagellin genes, and for the identification of isolates that may be able to undergo flagellum phase variation (Tominaga, 2004; Tominaga & Kutsukake, 2007).

As a preliminary validation of the EcOH database, we used it to screen 40 publically available *E. coli* genome assemblies that had reported O-groups. For 38 of these genomes the expected O-group (or O-cluster based on near-identical gene clusters (Iguchi *et al*., 2014)) was detected (**Table S2**). The two exceptions were *E. coli* isolates SE11 and SE15, which are reported in the literature as O152 and O150, respectively (Oshima *et al*., 2008; Toh *et al*., 2010). *In silico* analysis of these genomes identified *wzx* and *wzy* alleles for O16 and O173, respectively, and no BLAST+ hits to the alleles corresponding to the reported serogroups O152 and O150. Reported H-types were available for 20 of the reference genomes and we identified the expected H-alleles in all of these (**Table S2**), including both *fliC* H4 and *flnA* H17 in strain p12, consistent with a previous report of dual flagellin loci in this isolate (Ratiner *et al*., 2010).

### Comparison of serological phenotyping to in silico serotype prediction

We compared *in silico* serotyping (i.e., O- and H-genotyping) to serological phenotyping of 197 EPEC isolates. All isolates were serotyped by a national reference laboratory which yielded phenotypic identification of O-group for 144 isolates (73%; total 44 O-groups). The remaining 53 isolates were assessed either as O-rough (n=9, the isolates auto-agglutinated or were hyper-mucoid), or as O-non-typeable (n=44, agglutination did not occur with any antisera). H-types were phenotypically identified for 128 isolates (65% of those tested; 18 different H-types). The remaining 69 isolates were identified as H- (n=67, indicating that the isolate was non-motile) or H rough (n=2, indicating non-specific agglutination with H- antisera).

The 197 isolates were previously subjected to whole genome sequencing using the Illumina HiSeq platform (Ingle et al, 2015, in press). We compared two different strategies for *in silico* assignment of O- and H- types using the EcOH database: (i) typing direct from reads using SRST2, and (ii) *de novo* assembly (using Velvet Optimiser) followed by identification of alleles via BLAST+ (see **Methods**). Results are summarised in **Figure 1** with the full results reported in **Tables S3-S5**. SRST2 analysis yielded matching (i.e., same O-group) confident genotype calls at two O-determining loci (either *wzx* and *wzy*, or *wzm* and *wzt)* for 167 isolates (85%), and at one O-determining locus for a further 15 isolates. Thus, a total of 182 (93%) isolates were genotyped using SRST2, including 137/144 (95%) of those that were serologically typeable and 45/53 (85%) of those that were not (i.e., those that the reference laboratory identified as O-non-typeable or O-rough) (**Fig. 1(a)**). In comparison, BLAST+ analysis of the Velvet Optimiser assemblies yielded full-length (>90% coverage) hits to ≥1 O- gene locus for 180 (91%) isolates, including 135/144 (94%) of serologically typeable isolates and 45/53 (85%) of non-typeables (**Fig. 1(c)**). Alternative assemblies generated using SPAdes yielded fewer hits, with only 91/144 (64%) serologically typed isolates yielding full-length (>90% coverage) BLAST+ hits to any O-gene in the EcOH database. These assemblies were not analysed further.

**Figure 1.**
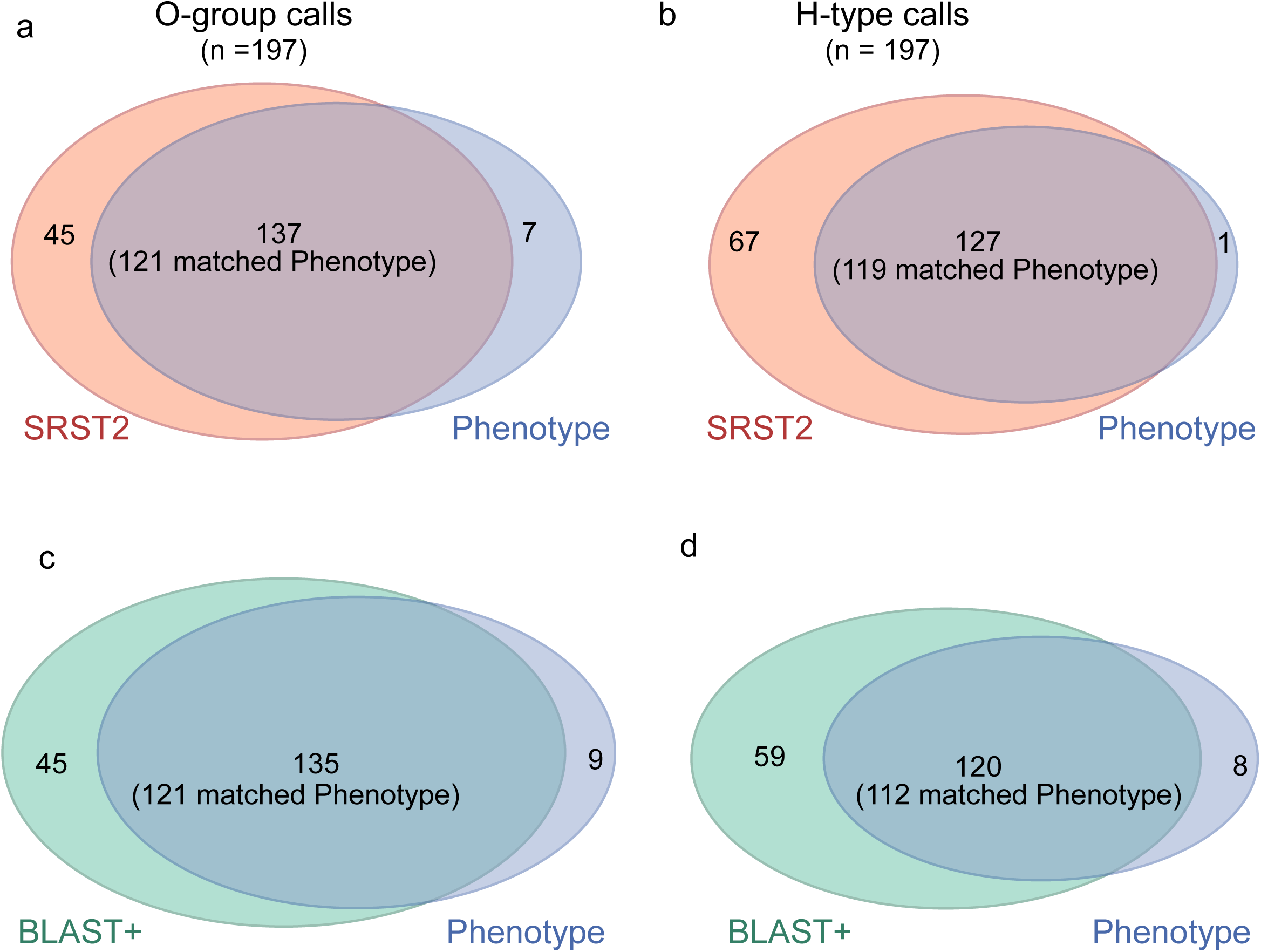
Comparison of serotype calls in 197 EPEC isolates. Venn diagrams showing the number of isolates yielding confident serotype calls for O-group or H-antigen by serological phenotyping *vs*. SRST2 analysis of reads (confident call at ≥1 O-antigen locus or at H locus; a-b), or *vs*. BLAST+ analysis of *de novo* assemblies (hit to ≥1 O-antigen locus or at H locus; c-d).

Of the 15 isolates for which SRST2 analysis did not yield a serotype call, 7 isolates had serological O-groups but no high confidence calls for any *wzx/wzy* or *wzm/wzt* genes (serologically typed as O2 [n=3], O103 [n=1], O108 [n=1], O124 [n=1] or O153 [n=1]). For 6 of 7 of these isolates, no O-antigen genes were detected in the assemblies either; for one isolate (serologically O103), SRST2 yielded a low-confidence call of O111, while assembly analysis detected O103 *wzx* and *wzy* alleles (**Table S3**). The remaining 8 isolates had no serotype detected via phenotypic or genotypic assays.

Of the 144 isolates that yielded a serological O-phenotype, genotyping based on confident SRST2 calls at ≥1 O-gene locus matched the serologically identified O-group for 121 isolates (84%), a different O-group for 16 isolates (11%; 15/16 with matching calls for both O loci) and no result for 7 isolates (5%) (**Table S3**). Of the 15 isolates for which SRST2 calls agreed at the two O loci but did not match the serological phenotype, assembly analysis identified the same O-group as SRST2 in 14/15 cases, and the serological O-type in 1/15 cases. There was only one isolate for which assembly-based analysis identified the same O-group as phenotyping when SRST2 had no result, and there were also two cases where SRST2 analysis identified the serological O-group and BLAST+ did not.

The possible reasons for mismatches between O-antigen phenotype and genotype include multiple genetic variants manifesting in the same phenotype (for example O45, see (Plainvert *et al*., 2007)) and/or atypical genetic variation such as multi-copy genes or novel genes. To explore these possibilities we manually inspected the genome assemblies of isolates yielding conflicting genotype/phenotype calls and identified twelve novel O-antigen loci, which were added to the EcOH database with the suffix ‘var1, var2’ etc., to differentiate them from prototypical alleles. (**Fig. S1**). For example, three isolates phenotyped as O116 or O33-related, had detectable *wzx* O116 genes but no *wzy* genes. Interestingly, the *wzx* alleles were detected at high depth (~100x) and were highly divergent from the reference O116 *wzx* allele (~10% nucleotide divergence, the maximum limit of detection we used for genotyping). We hypothesised that these isolates may carry *wzy* genes that are genetically distant from the prototypical alleles that were included in our database, but which nonetheless result in similar phenotypic agglutination patterns to isolates carrying more prototypical genes. Investigation of the corresponding genome assemblies revealed a novel O-antigen locus, including novel *wzx* and *wzy* variants that were 10% and 40% divergent, respectively, from the prototypical O116 gene sequences (**Fig. S1(a)**). The novel *wzx* and *wzy* sequences were labelled O116var1 and added to the EcOH database to facilitate identification of this novel type in future. In the genomes of four isolates genotyped as O8 but phenotyped as O153 (**Table S3**), we confirmed the presence of O8 *wzx* and *wzy* alleles, but also identified a putative *wzx* homolog (with 76% identity to O83) outside the O-antigen region which, if expressed, could potentially alter the O-antigen phenotype (**Fig. S1(b)**). Interestingly, an isolate which displayed the O153 phenotype but had no O-locus hits in the EcOH database, also had two putative novel *wzx* genes (53% nucleotide similarity to O83 and 57% to O170, respectively), one of which was similar to the additional *wzx* gene identified in the genomes of the O153 phenotype/O8 genotype isolates (**Fig. S1(b)**).

H-typing using the EcOH database yielded similar results to O-typing. SRST2 analysis returned 127 confident calls that matched the phenotype in 119 of 128 (93%) of the serologically H-typed isolates, and gave confident genotype calls in 67 of 69 (97%) non-motile (serologically H-) isolates (**Fig. 1(b)**, **Tables S4–5**). In contrast, assembly analysis identified the expected genes in only 112/128 (88%) of serologically H-typed isolates and 59/69 (86%) of non-motile isolates (**Fig. 1(d)**, **Tables S4–5**). The high rate of genotype calls amongst phenotypically H-non-motile isolates is likely due to a lack of flagellin expression during serotyping, which does not affect genotyping. Only two isolates had no flagellin genes detected from the sequence data. These were non-motile and may be the only isolates that genuinely lack the ability to express flagella.

### Applications of rapid in silico serotyping and multi-locus sequence typing of E. coli

The data above show that the use of SRST2 and the EcOH database to type raw Illumina read sets can provide rapid *in silico* serotyping, that outperforms assembly-dependent analysis (especially for H-typing) and is largely predictive of results obtained from serological typing while yielding fewer ‘untypeable’ results. In addition to the EcOH database, other databases, such as those used for multi-locus sequence typing (MLST) and antibiotic resistance gene profiles, can be interrogated using SRST2 (Inouye *et al*., 2014), with a single SRST2 analysis returning MLST, serotype and antimicrobial resistance gene results in a few minutes (approximately 5–10 minutes for paired Illumina data at mean read depth 50-100x, see (Inouye *et al*., 2014)). We therefore sought to demonstrate the utility of this approach for the rapid characterisation of *E. coli* genomes, including serotyping, MLST and antibiotic resistance gene profiling, in a variety of contexts including the investigation of EPEC and ETEC pathotype populations, the epidemic UPEC clone ST131, and isolates associated with foodborne outbreaks.

SRST2 analysis of the 197 EPEC plus 360 recently sequenced ETEC isolates (Mentzer *et al*., 2014) highlighted that both pathotypes comprise a diversity of phylogenetic lineages and serotypes (**Fig. 2**, **Fig. S2**). A total of 46 O- and 20 H-types amongst the EPEC isolates; and 54 O- and 31 H-types amongst the ETEC isolates (**Fig. 2**, **Fig. S2**). These analyses suggest that H-types are more stably maintained within *E. coli* clones than are O-groups (**Fig. 3)**, consistent with observations of serological diversity in *Salmonella enterica* (Achtman *et al*., 2012). Interestingly, some MLST sequence types (STs) showed greater evidence of O-group diversity than others (**Figs. 2–3, Fig S2**), particularly ST29 and ST517 (EPEC); ST155, ST 173 and ST23 (ETEC); and ST10 (both EPEC and ETEC). ST10 was frequent amongst both EPEC and ETEC and displayed high O-group diversity in both pathotypes (Simpson index = 0.62 in EPEC, 0.84 in ETEC; see **Fig. 3**). Interestingly, all ST10 EPEC carried H40 flagella, but ST10 ETEC had 8 different H-types (Simpson index = 0.79; see **Fig. 3**). This high diversity within ST10 is consistent with the fact that it was one of the first *E. coli* lineages identified as harbouring multiple pathotypes as well as commensal strains (Mentzer *et al*., 2014; Wirth *et al*., 2006).

**Figure 2.**
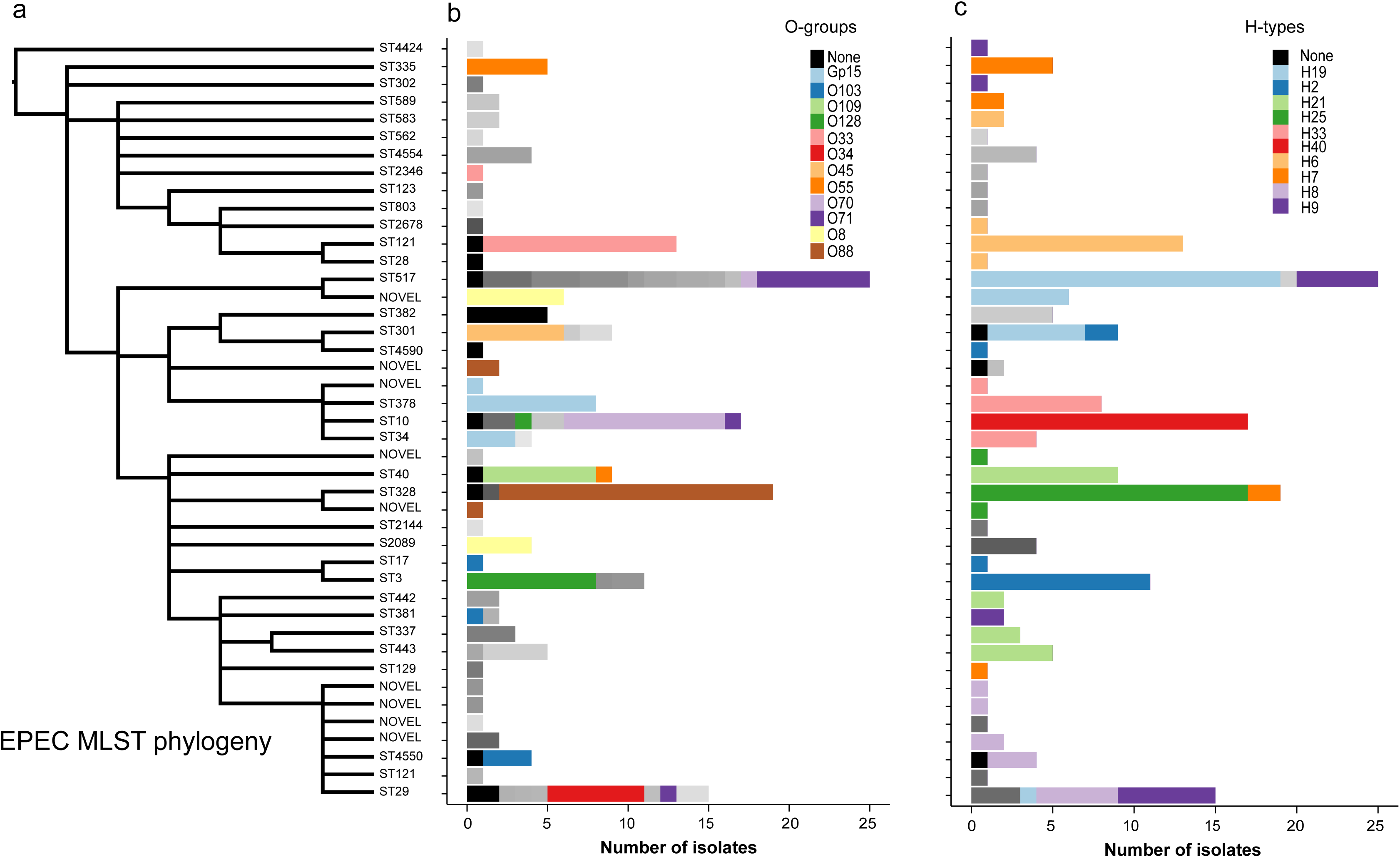
Serotype and sequence type diversity amongst 197 EPEC isolates. **(a)** Tree of EPEC isolates based on clustering of MLST allele profiles. Tips are labelled with sequence types. **(b)** Distribution of O-groups determined by SRST2 analysis using the EcOH database. The 12 most frequent O-groups are highlighted in colour with the less frequent O-groups shown in grey. **(c)** Distribution of H-types detected by SRST2 analysis using the EcOH database. The 10 most frequent H-types are highlighted in colour with the less frequent H-types shown in grey.

**Figure 3.**
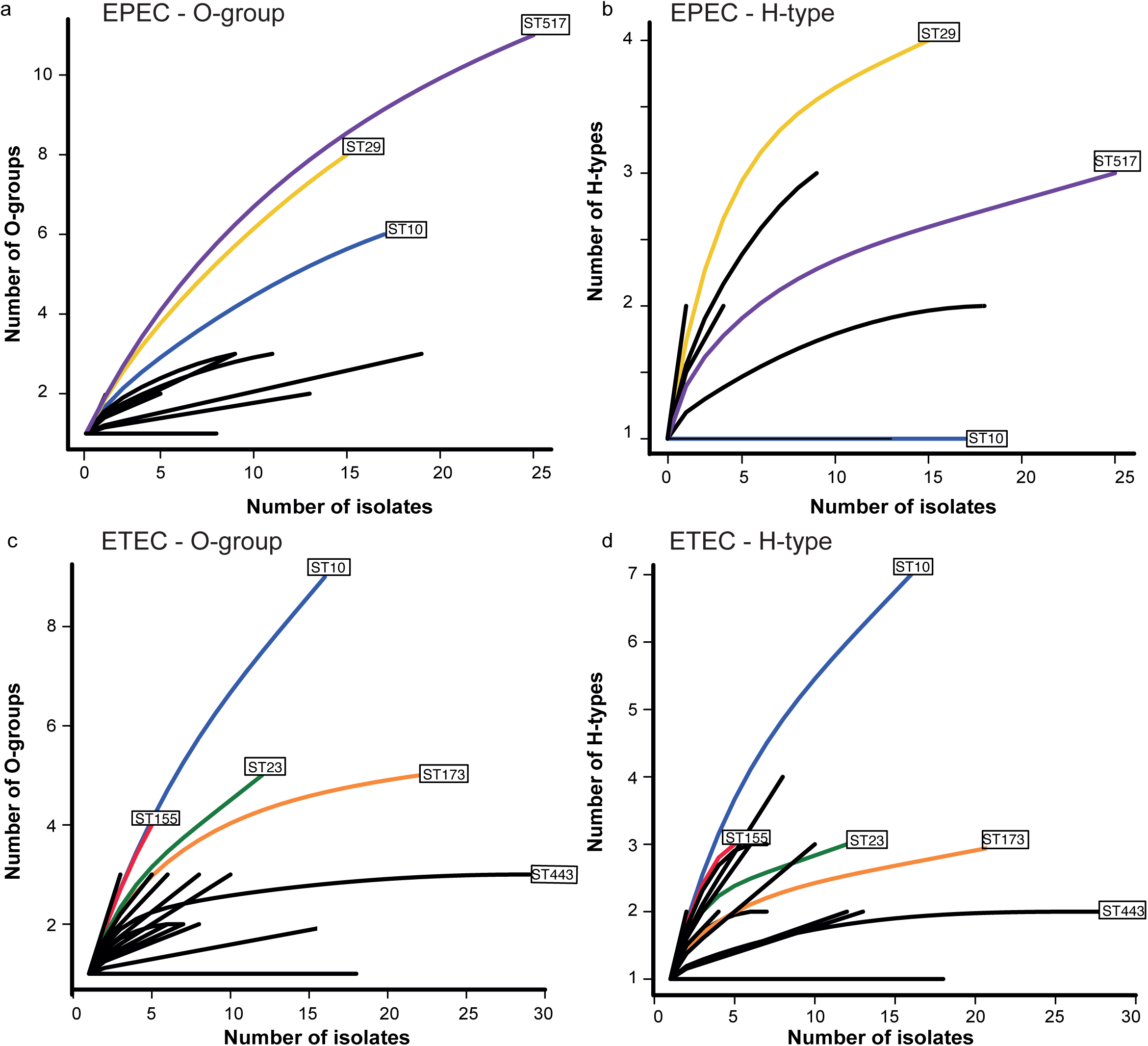
Accumulation curves for O-groups and H-types amongst EPEC and ETEC isolates. Accumulation of O-groups (**a, c**) or H-types (**b, d**) detected as more isolates in each lineage (ST) of EPEC (a, b) or ETEC (c, d) are sampled.

Next we used SRST2 and the EcOH database to analyse Illumina read sets for 169 UPEC isolates previously reported as belonging to the epidemic UPEC clone ST131 (Petty *et al*., 2014; Price *et al*., 2013) (**Fig. 4)**. Most isolates were confirmed as ST131, although 6 were single locus variants of ST131, including four belonging to ST95 (consistent with the original report on these genomes (Petty *et al*., 2014)). *In silico* serotyping identified the majority of isolates (90%) as O25:H4, which is the reported O-group for this epidemic clone (Nicolas-Chanoine *et al*., 2008). However, we also identified 14 isolates (8%) as O16:H5; these clustered together tightly in the core genome phylogeny, indicating they represent a subclone of ST131 in which a change of serotype has occurred (**Fig. 4)**. The O16:H5 sub-clone carried fewer resistance genes than other ST131 genomes and corresponds to ST131 Clade A, which has been identified as an ancestral sub-lineage of ST131 that is distinct from the sub-lineage which is now globally disseminated (Petty *et al*., 2014). O-antigen variation within ST131 was detected in the original genome reports (Petty *et al*., 2014; Price *et al*., 2013), but was not explored in detail. Our data highlight the utility of *in silico* serotyping to illuminate on-going microevolution in important epidemic clones of *E. coli*, including change of serotype, which could confound serological identification of outbreak-related isolates.

**Figure 4.**
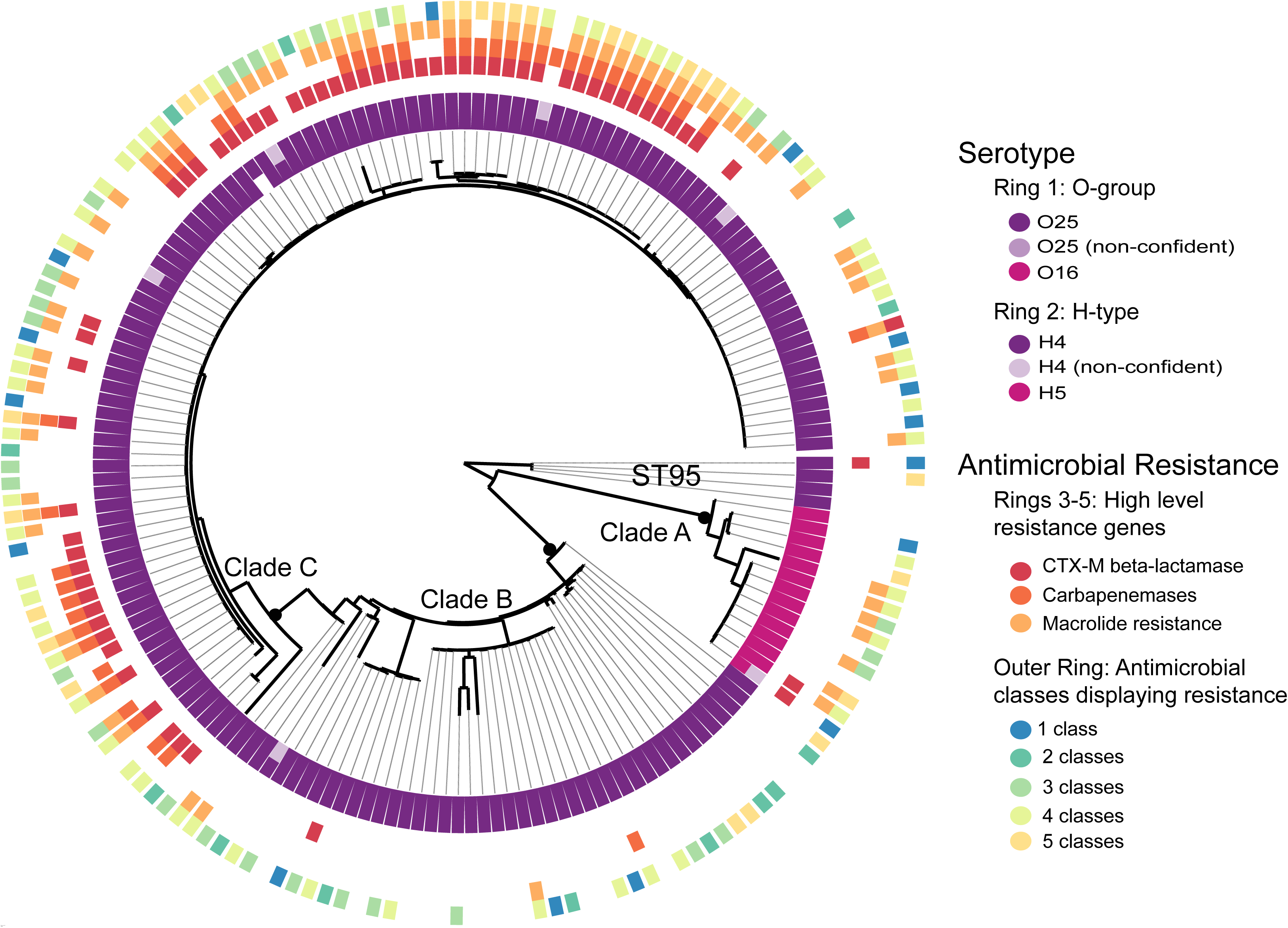
*In silico* prediction of serotype and antimicrobial resistance for UPEC ST131 isolates. Core genome SNP tree for 170 isolates, outgroup-rooted using ST95 isolates. Serotype and acquired antimicrobial resistance gene profiles, detected using SRST2, are demarcated on the rings surrounding the phylogeny; note low-confidence serotype calls are shown in paler colours.

Finally, we performed *in silico* serotyping of 300 foodborne outbreak-associated *E. coli* genomes recently deposited by public health laboratories into the GenomeTrakr project (NCBI BioProject accession: PRJNA183844). **Figure 5** shows our *in silico* serotyping results overlaid on the GenomeTrakr kmer-based tree. Environmental isolates displayed a diversity of ST, O- and H-types, whereas most clinical isolates belonged to one of six clonal lineages, each characterised by a specific serotype (**Fig. 5)**. The predominant lineage amongst these recently-deposited isolates was the well characterised enterohaemorrhagic *E. coli* (EHEC) lineage ST11, O157:H7. Other lineages included ST16 O111:H8, ST655 O121:H19, and ST232 O145:H- (in which no serotype variation was detected), as well as clonal complexes (CC) CC21 O26:H11 and CC17 O103:H2 (both of which displayed some serotype variation, see **Fig. 5)**. For 227 isolates (76%), matching confident calls were obtained for both O-antigen genes, whilst 272 isolates (91%) had a confident call for at least one allele. In most cases where a low-confidence O-antigen genotype call was made (due to low read depth), the call was for O157 alleles; the position of these genomes within the ST11 O157:H7 lineage of the kmer tree suggests that these low-confidence calls of these genomes are likely to be correct. Only 7 isolates (2%) yielded no genotype calls for the H-locus, indicating they are likely to be non-motile. These data demonstrate the utility of our method for *in silico* serotype prediction of *E. coli* sequenced for the investigation of foodborne outbreaks or other purposes.

**Figure 5.**
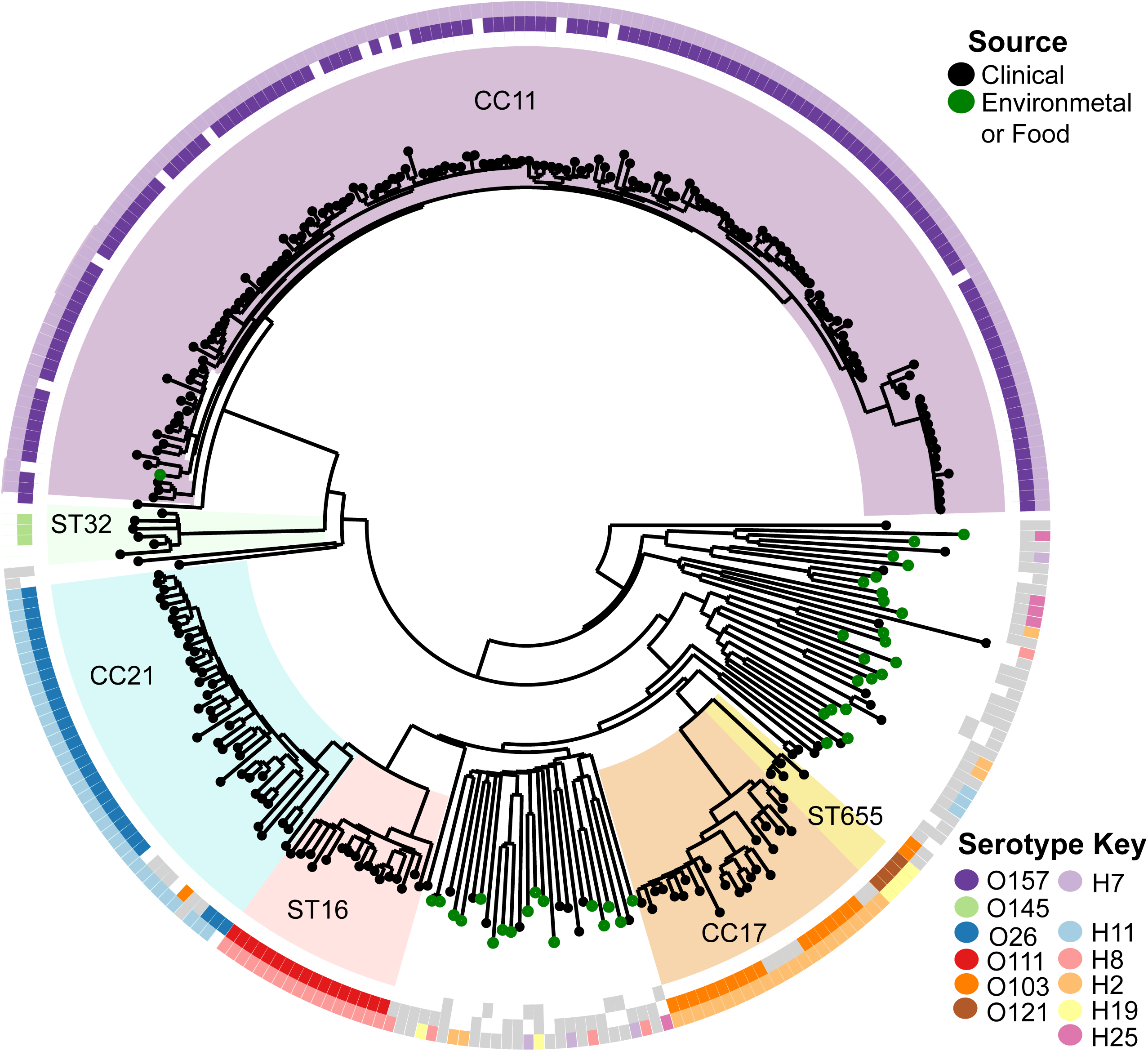
*In silico* prediction of serotype and sequence type from GenomeTrakr data. Kmer-based tree obtained from the GenomeTrakr project; common sequence types (STs) or clonal complexes (CC), identified by SRST2, are highlighted and labelled. Rings show predicted O-type (inner ring) and H-type (outer ring).

## Conclusion

This study has demonstrated that *E. coli* O- and H-genotypes can be rapidly and accurately extracted from whole genome data using the free SRST2 software and a new publicly available database, EcOH. The method improves on both (i) serological phenotyping, which is resource intensive in terms of time, labour and reagent costs, and fails to type up to one third of *E. coli* isolates, and (ii) assembly-based approaches for *in silico* genotyping of Illumina data, particularly for H-typing, which are more computationally expensive and are highly dependent on the quality of the sequence data. Importantly, since SRST2 works on raw reads and can readily be used to extract other useful genotyping information in addition to serotype, including MLST, antimicrobial resistance and virulence genes (Inouye *et al*., 2014), it lends itself to integration with robust assembly-free pathogen genome characterisation pipelines. Our data demonstrate that this approach can be used to readily infer serotypes from genome data currently being produced by GenomeTrakr and other public health networks as part of routine investigation of foodborne *E. coli* outbreaks. This could be useful in identifying the emergence of novel serotypes within outbreak clades that may signal a shift in the pathogen population during its dissemination (as we identified in ST131 UPEC), and importantly will provide backwards compatibility with the wealth of serotype data that is currently available from historical outbreak investigations.

## Acknowledgements

This work was supported by the Australia NHMRC (Project Grants #1043830 to KEH, #1009296 and #1067428 to RRB; Fellowship #1061409 to KEH; Fellowship #1061435 to MI (co-funded by the Australian Heart Foundation Career Development Fellowship)); the Bill & Melinda Gates Foundation (Grant #38874 to MML); and the Victorian Life Sciences Computation Initiative (VLSCI) (Grant #VR0082). We thank Gordon Dougan and the sequencing teams at the Wellcome Trust Sanger Institute for sequencing the EPEC isolate collection.

**Abbreviations**
CC: clonal complex
EHEC: enterohaemorrhagic *Escherichia coli*
EPEC: enteropathogenic *Escherichia coli*
ETEC: enterotoxigenic *Escherichia coli*
ML: Maximum Likelihood
MLST: multi-locus sequence typing
SNP: single nucleotide polymorphism
UPEC: uropathogenic *Escherichia coli*

## References

Achtman, M., Wain, J., Weill, F.-X., Nair, S., Zhou, Z., Sangal, V., Krauland, M. G., Hale, J. L., Harbottle, H. & other authors. (2012). Multilocus Sequence Typing as a Replacement for Serotyping in Salmonella enterica. PLoS Pathog 8, e1002776–19.

Bankevich, A., Nurk, S., Antipov, D., Gurevich, A. A., Dvorkin, M., Kulikov, A. S., Lesin, V. M., Nikolenko, S. I., Pham, S. & other authors. (2012). SPAdes: A New Genome Assembly Algorithm and Its Applications to Single-Cell Sequencing. Journal of Computational Biology 19, 455–477.

Bao, E., Jiang, T. & Girke, T. (2014). AlignGraph: algorithm for secondary de novo genome assembly guided by closely related references. Bioinformatics 30, i319–i328.

Boetzer, M., Henkel, C. V., Jansen, H. J., Butler, D. & Pirovano, W. (2011). Scaffolding pre-assembled contigs using SSPACE. Bioinformatics 27, 578–579.

Boetzer, M. & Pirovano, W. (2012). Toward almost closed genomes with GapFiller. Genome Biol 13, R56.

Carver, T., Berriman, M., Tivey, A., Patel, C., Bohme, U., Barrell, B. G., Parkhill, J. & Rajandream, M. A. (2008). Artemis and ACT: viewing, annotating and comparing sequences stored in a relational database. Bioinformatics 24, 2672–2676.

Chandler, M. E. & Bettelheim, K. A. (1974). A rapid method of identifying *Escherichia coli* H antigens. Zentralblatt für Bakteriologie, Parasitenkunde, Infektionskrankheiten und Hygiene Abt, Originale 229, 74–79.

Croxen, M. A., Law, R. J., Scholz, R., Keeney, K. M., Wlodarska, M. & Finlay, B. B. (2013). Recent advances in understanding enteric pathogenic *Escherichia coli*. Clin Microbiol Rev 26, 822–880.

DebRoy, C., Roberts, E. & Fratamico, P. M. (2011). Detection of O antigens in *Escherichia coli*. Anim Health Res Rev 12, 169–185.

Feng, L., Liu, B., Liu, Y., Ratiner, Y. A., Hu, B., Li, D., Zong, X., Xiong, W. & Wang, L. (2008). A genomic islet mediates flagellar phase variation in *Escherichia coli* strains carrying the flagellin-specifying locus *flk*. J Bacteriol 190, 4470–4477.

Feng, L., Senchenkova, S. N., Yang, J., Shashkov, A. S., Tao, J., Guo, H., Cheng, J., Ren, Y., Knirel, Y. A. & other authors. (2004). Synthesis of the heteropolysaccharide O Antigen of *Escherichia coli* O52 requires an ABC transporter: structural and genetic evidence. J Bacteriol 186, 4510–4519.

Fratamico, P. M., DebRoy, C., Miyamoto, T. & Liu, Y. (2009). PCR detection of enterohemorrhagic *Escherichia coli* O145 in food by targeting genes in the *E. coli* O145 O-antigen gene cluster and the Shiga Toxin 1 and Shiga Toxin 2 genes. Foodborne Pathogens and Disease 6, 605–611.

Gupta, S. K., Padmanabhan, B. R., Diene, S. M., Lopez-Rojas, R., Kempf, M., Landraud, L. & Rolain, J. M. (2013). ARG-ANNOT, a new bioinformatic tool to discover antibiotic resistance genes in bacterial genomes. Antimicrobial Agents and Chemotherapy 58, 212–220.

Harmon, L. J., Weir, J. T., Brock, C. D., Glor, R. E. & Challenger, W. (2007). GEIGER: investigating evolutionary radiations. Bioinformatics 24, 129–131.

Iguchi, A., Iyoda, S., Kikuchi, T., Ogura, Y., Katsura, K., Ohnishi, M., Hayashi, T. & Thomson, N. R. (2014). A complete view of the genetic diversity of the *Escherichia coli* O-antigen biosynthesis gene cluster. DNA Res.

Ingle, D.J., Tauschek, M., Edwards, D.J., Hocking, D.M., Pickard, D.J., Azzopardi, K.I., Amarasena, T., Bennett-Wood, V., Pearson, J.S., & other authors. Evolution of atypical enteropathogenic E. coli by repeated acquisition of LEE pathogenicity island variants *Nat Micro* In press

Inouye, M., Dashnow, H., Raven, L., Schultz, M. B., Pope, B. J., Tomita, T., Zobel, J. & Holt, K. E. (2014). SRST2: Rapid genomic surveillance for public health and hospital microbiology labs. Genome Med 6, 1–16.

Jenkins, C. (2015). Whole-Genome sequencing data for serotyping *Escherichia coli* - It’s time for a change! J Clin Microbiol 53, 2402–2403.

Joensen, K. G., Tetzschner, A. M. M., Iguchi, A., Aarestrup, F. M. & Scheutz, F. (2015). Rapid and easy *in silico* serotyping of *Escherichia coli* using whole genome sequencing (WGS) data. J Clin Microbiol 2410–2426.

Kauffmann, F. (1944). The basis of O Antigen Serotyping is the standard tube agglutination tests adapted to use U-bottomed microtitre trays. Microbiologica Scandinavia 21, 20–40.

Krogh, A., Larsson, B., Heijne, von, G. & Sonnhammer, E. L. L. (2001). Predicting transmembrane protein topology with a hidden markov model: application to complete genomes. J of Mol Biol 305, 567–580.

Kwong, J. C., McCallum, N., Sintchenko, V. & Howden, B. P. (2015). Whole genome sequencing in clinical and public health microbiology. Pathology 47, 199–210.

Langmead, B., Trapnell, C., Pop, M. & Salzberg, S. L. (2009). Ultrafast and memory-efficient alignment of short DNA sequences to the human genome. Genome Biol 10, R25.

Li, D., Bin Liu, Chen, M., Guo, D., Guo, X., Liu, F., Feng, L. & Wang, L. (2010). A multiplex PCR method to detect 14 *Escherichia coli* serogroups associated with urinary tract infections. Journal of Microbiological Methods 82, 71–77.

Li, H., Handsaker, B., Wysoker, A., Fennell, T., Ruan, J., Homer, N., Marth, G., Abecasis, G., Durbin, R.1000 Genome Project Data Processing Subgroup. (2009). The Sequence Alignment/Map format and SAMtools. Bioinformatics 25, 2078–2079.

Liu, D., Cole, R. A. & Reeves, P. R. (1996). An O-antigen processing function for Wzx(RfbX): a promising candidate for O-unit flippase. J Bacteriol 178, 2102–2107.

Mentzer, von, A., Connor, T. R., Wieler, L. H., Semmler, T., Iguchi, A., Thomson, N. R., Rasko, D. A., Joffre, E., Corander, J. & other authors. (2014). Identification of enterotoxigenic *Escherichia coli* (ETEC) clades with long-term global distribution. Nat Genet 46, 1321–1326.

Nicolas-Chanoine, M.-H., Blanco, J., Leflon-Guibout, V., Demarty, R., Alonso, M. P., Caniça, M. M., Park, Y.-J., Lavigne, J.-P., Pitout, J. & Johnson, J. R. (2008). Intercontinental emergence of Escherichia coli clone O25:H4-ST131 producing CTX-M-15. J Antimicrob Chemother 61, 273–281.

Oksanen, J., Blanchet, F. G., Kindt, R., Legendre, P., Minchin, P. R., OHara, R. B., Simpson, G. L., Solymos, P., Stevens, M. H. H. & Wagner, H. (2015). vegan: Community Ecology Package.

Oshima, K., Toh, H., Ogura, Y., Sasamoto, H., Morita, H., Park, S. H., Ooka, T., Iyoda, S., Taylor, T. D. & other authors. (2008). Complete genome sequence and comparative analysis of the wild-type commensal *Escherichia coli* strain SE11 isolated from a healthy adult. DNA Res 15, 375–386.

Paradis, E., Claude, J. & Strimmer, K. (2004). APE: Analyses of Phylogenetics and Evolution in R language. Bioinformatics 20, 289–290. Oxford University Press.

Petty, N. K., Ben Zakour, N. L., Stanton-Cook, M., Skippington, E., Totsika, M., Forde, B. M., Phan, M. D., Gomes Moriel, D., Peters, K. M. & other authors. (2014). Global dissemination of a multidrug resistant *Escherichia coli* clone. Proc Natl Acad Sci USA 111,5694–5699.

Plainvert, C., Bidet, P., Peigne, C., Barbe, V., Medigue, C., Denamur, E., Bingen, E. & Bonacorsi, S. (2007). A new O-antigen gene cluster has a key role in the virulence of the *Escherichia coli* Meningitis clone O45:K1:H7. J Bacteriol 189, 8528–8536.

Price, L. B., Johnson, J. R., Aziz, M., Clabots, C., Johnston, B., Tchesnokova, V., Nordstrom, L., Billig, M., Chattopadhyay, S. & other authors. (2013). The epidemic of Extended-Spectrum-ß-Lactamase-Producing *Escherichia coli* ST131 is driven by a single highly pathogenic subclone, *H30*-Rx. mBio 4, 1–10.

Ratiner, Y. A. (1998). New Flagellin-Specifying Genes in some *Escherichia colis* strains. J Bacteriol 180, 979–984.

Ratiner, Y. A., Sihvonen, L. M., Liu, Y., Wang, L. & Siitonen, A. (2010). Alteration of flagellar phenotype of *Escherichia coli* strain P12b, the standard type strain for flagellar antigen H17, possessing a new *non-fliC* flagellin gene *flnA*, and possible loss of original flagellar phenotype and genotype in the course of subculturing through semisolid media. Arch Microbiol 192, 267–278.

Rice, P., Longden, I. & Bleasby, A. (2000). EMBOSS: The European Molecular Biology Open Software Suite. Trends Genet 16, 276–277.

Samuel, G. & Reeves, P. (2003). Biosynthesis of O-antigens: genes and pathways involved in nucleotide sugar precursor synthesis and O-antigen assembly. Carbohydrate Research 338, 2503–2519.

Seemann, T. (2014). Prokka: rapid prokaryotic genome annotation. Bioinformatics 30, 2068–2069.

Toh, H., Oshima, K., Toyoda, A., Ogura, Y., Ooka, T., Sasamoto, H., Park, S. H., Iyoda, S., Kurokawa, K. & other authors. (2010). Complete Genome Sequence of the Wild-Type Commensal Escherichia coli Strain SE15, Belonging to Phylogenetic Group B2. J Bacteriol 192, 1165–1166.

Tominaga, A. (2004). Characterization of six flagellin genes in the H3, H53 and H54 standard strains of *Escherichia coli*. Genes Genet Syst 79, 1–8.

Tominaga, A. & Kutsukake, K. (2007). Expressed and cryptic flagellin genes in the H44 and H55 type strains of *Escherichia coli*. Genes Genet Syst 82, 1–8.

Wang, L., Rothemund, D., Curd, H. & Reeves, P. R. (2003). Species-wide variation in the *Escherichia coli* flagellin (H-antigen) gene. J Bacteriol 185, 2936–2943.

Wattam, A. R., Abraham, D., Dalay, O., Disz, T. L., Driscoll, T., Gabbard, J. L., Gillespie, J. J., Gough, R., Hix, D. & other authors. (2013). PATRIC, the bacterial bioinformatics database and analysis resource. Nucleic Acids Res 42, D581–D591.

Wirth, T., Falush, D., Lan, R., Colles, F., Mensa, P., Wieler, L. H., Karch, H., Reeves, P. R., Maiden, M. C. J. & other authors. (2006). Sex and virulence in *Escherichia coli:* an evolutionary perspective. Mol Microbiol 60, 1136–1151.

Zerbino, D. R. & Birney, E. (2008). Velvet: Algorithms for de novo short read assembly using de Bruijn graphs. Genome Res 18, 821–829.

